# Sequencing-based Spatial Transcriptomics with scRNA-seq Sensitivity

**DOI:** 10.1101/2025.01.15.633111

**Authors:** Renjie Liao, Defeng Fu, Zaoxu Xu, Han Liang, Xiaoran Zhou, Yiling Chen, Xueqi Liu, Jiajun Cheng, Ruidong Guo, Chen Li, Huihua Xia, Gailing Li, Diewen Feng, Wei Chen, Yang Chen, Longchao Chen, Zihang Huang, Yang Zhou, Qingbin Chen, Dexia Chen, Shuguang Xuan, Changling Zhang, Yongyi Lu, Hui Wang, Taiqing Feng, Yuanye Bao, Luyang Zhao, Erkai Liu, Gufeng Wang

## Abstract

The advent of spatial transcriptomics has dramatically expanded our ability to study the vast network of cell-cell interactions at the molecular level in tissue. Among current methods, sequencing-based approaches have great potential in discovering because of its unbiased capture. In the last couple of years, the spatial resolution for the capture addresses has been significantly improved from 100 μm to <1 μm, well below the size of a mammalian cell. However, the capture efficiency has always been a pain point, ∼one order of magnitude lower than that of single cell RNA sequencing (scRNA-seq). The low capture efficiency limits the depth and breadth of its applications in the study of complex biological systems and diseases. Here, we introduce Salus Spatial transcriptomic system (Salus-STS), which provides ∼1 μm capture resolution and a capture efficiency ∼1 order of magnitude higher than other current methods. Analysis with sub-cellular resolution becomes practical for sequencing-based spatial transcriptomics.

---

It has been long ever since researchers started to explore gene expression using RNA transcripts at cell- and tissue-levels^1^. Initial efforts included those hybridization probe-based imaging methods, like fluorescence in situ hybridization (FISH)^2^. Not until very recently can this approach be called spatial transcript-OMICS (e.g., MERFISH^3^, seqFISH^4^ and STARmap^5^) when a large panel of genes (over several hundred) can be studied in one experiment. However, these methods still rely on preliminary knowledge and well-designed panels, limiting their roles in discovering.

Sequencing-based spatial transcriptomics methods, on the other hands, capture the whole genome expression profile using spatially barcoded probes. For this reason, these techniques are also referred to as “unbiased methods”. A group of techniques emerged, such as Visium^6^, Slide-seq^7,8^, HDST^9^, DBIT^10^, Decoder-Seq^11^, Seq-Scope^12^, Stereo-seq^13^, Open-ST^14^, Nova-ST^15^, PixelSeq^16^, Visium HD^17^, which drastically expedited the advance of relevant fields. So far, they have shown their ability to determine cell-type architecture of tissue, reveal cell–cell interactions, and monitor cell differentiation and tissue development, etc.^18–20^ Some of these methods’ spatial resolution for capture addresses is improved to be 0.5∼2 μm^21^, sufficiently high to support cell-level studies.

However, current sequencing-based methods exhibit poor capture efficiency of mRNA molecules. While a mammalian cell typically has 10^5^–10^6^ mRNA molecules, and up to 10,000 different gene expressed^22^, spatial transcriptomics experiments usually report the detection of mRNAs in lower hundreds per 10 × 10 μm² area on a capture chip, a size similar to that of a cell.

## Compared to single cell RNA sequencing that can capture several thousand mRNA molecules^23^, the capture efficiency of spatial transcriptomics is one order of magnitude lower

An accurate gene expression profile is essential for subsequent bioinformatics analysis. The greater the number and variety of transcripts that can be detected, the more comprehensive and accurate the transcriptomic state of a tissue is. Low capture efficiency has several major implications: (1) difficulty in cell type identification and classification; (2) missing rare cell types and sub-types; (3) missing low-abundance genes that may play a key role in biological processes; (4) inaccurate cell-cell interactions due to missing expressed genes; (5) limited resolving power for dynamic gene expression changes upon external perturbation; (6) susceptible to background noises such as non-specifical binding, sequencing error, etc. In this manuscript, we designed a new spatial transcriptomic system (Salus-STS), which provides ∼1 μm capture resolution and a much-enhanced capture efficiency.

## RESULTS

### Capture substrate design

Consider an equilibrium surface binding reaction: *S* + *A* = *SA*, where *S* stands for the surface binding site, *A* stands for the target molecule being captured, *SA* stands for the target-surface site complex, the surface concentration of captured targets can be approximated as [*SA*] = *cK*/(*1*+*cK*)*[*S0*] when the depletion of the reactant *A* is negligible, where [*SA*] and [*S0*] stand for the surface concentration of reacted and initial binding sites, respectively, *c* is the initial concentration of reactant *A* in the solution, and *K* is the binding constant. Note that it is assumed that the releasing of mRNAs from the tissue is complete. In this case, when the depletion of mRNA is on ∼1% level or smaller, our early negligible depletion assumption becomes valid. It is clear that the amount of the mRNAs captured is directly proportional to the initial surface concentration of the capture sites. Thus, the key to improve mRNA capture efficiency is to increase the amount of capture probes per unit area.

Bearing this in mind, we designed Salus-STS capture substrate as depicted in Fig. 1. A next generation sequencer Salus Pro^TM^ by Salus BioMed operating under the sequencing-by-synthesis (SBS) principle^24,25^ served as the platform for generating ultra-high-density capture probes on substrate with random spatial barcodes (SBCs), and for decoding the SBCs. Note that there are recent efforts using Illumina sequencing flowcells to fabricate the capture substrates, e.g., Seq-Scope^12^, Open-ST^14^, Nova-ST^15^, etc. However, since they have to use the P5 and P7 primers on the flowcell surface, their capability in optimizing the experiments is largely hindered.

**Figure 1.**
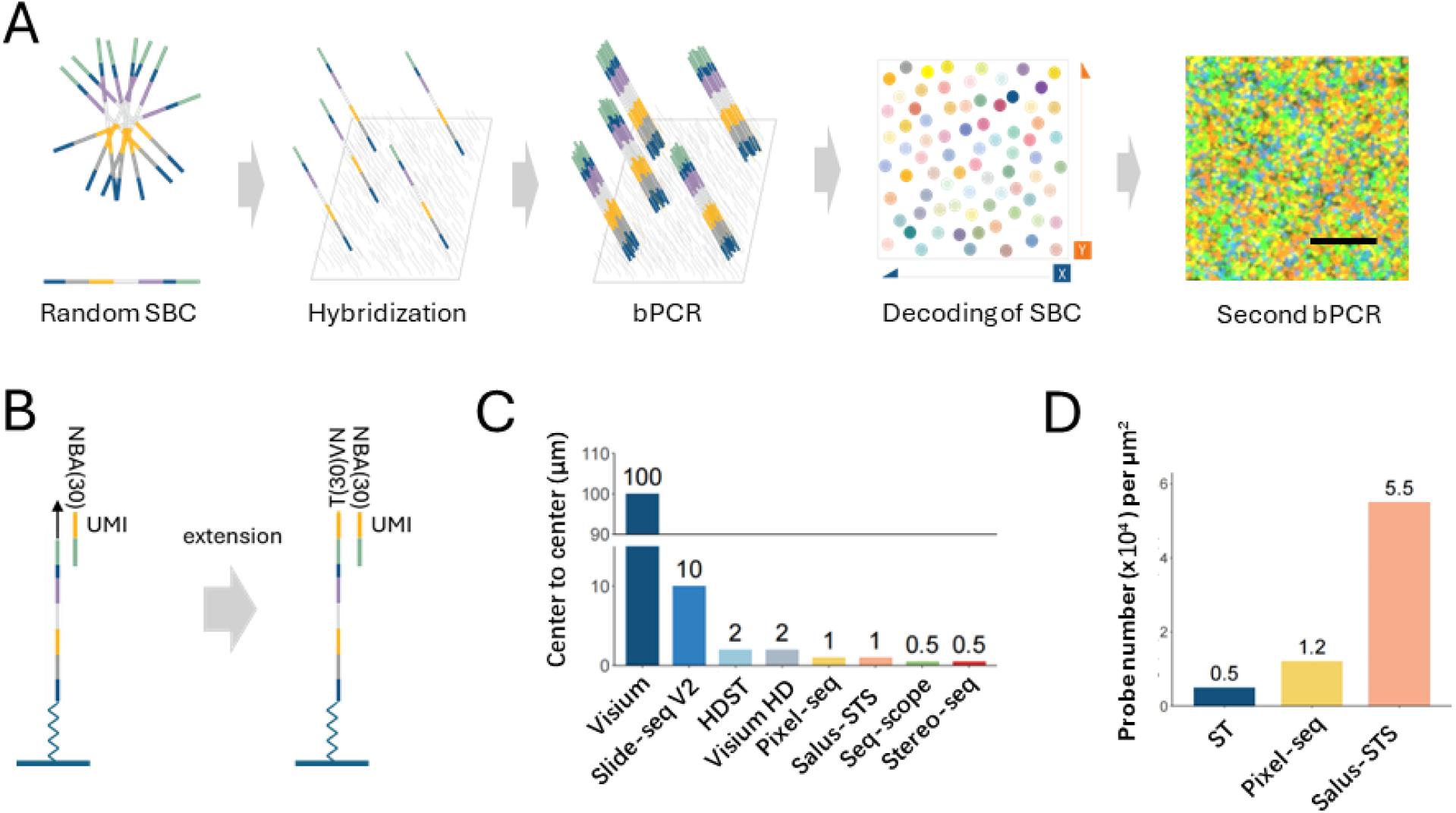
Salus-STS capture substrate design and characterization. (A) Schematics illustrating the generation of high-density clusters of the stalk portion of the capture probes. Scale bar: 10 μm. SBC: Spatial barcodes; bPCR: bridge polymerase chain reaction. (B) Attachment of molecular recognition portion to capture probe stalks. (C) Comparison of center-to-center distance of different methods in the literature. (D) Comparison of reported probe densities.

Here, we designed the capture substrate so that it is compatible with a Salus Pro^TM^ DNA sequencer. The capture areas are either 8.6×8.6 mm^2^ or 11×11 mm^2^. The primers on the surface for bridge amplification were redesigned so they would not interfere with standard Illumina library construction. The stalk portion of the capture probes contained a random 30-bp nucleotide sequence serving as the spatial barcode, which holds excess complexity (over 10^18^) for spatial mapping over an area larger than 1 cm^2^.

In the fabrication of the capture substrate, the stalks of the capture probes were bridge amplified on the substrate with optimized primer surface density. Through multiple rounds of amplification, clusters of the stalks, each with ∼10,000 copies, formed with a surface density of ∼1 cluster/μm^2^, equivalent to a center-to-center distance of ∼1 μm (Fig. 1A). In this setup, the stalks of the capture probes were randomly hybridized on the surface, leading to randomly distributed probes. Subsequently, the Salus Pro^TM^ sequencer was employed to decode random SBCs for each physical address on the substrate.

Following decoding, the molecular recognition portions of the probes were added to the stalks with a guided extension approach (Fig. 1B). These recognition portions consist of unique molecular identifiers (UMIs) and polyT sequences. UMIs are random 10-bp nucleotides serving to distinguish transcripts of the same gene locus captured by the same cluster of probes. PolyT sequences were used to capture mRNAs in a unbiased manner through hybridization with mRNAs’ polyA tails.

Salus-STS has a spatial capture resolution of ∼1 μm (Fig. 1C), reaching the highest tier among current spatial transcriptomic technologies. To increase the number of probes on the surface, there are two ways: (1) increase the amplification rounds in the first bPCR step, or (2) add an additional bPCR step (Fig. 1A) after decoding. The latter can fill up all the open surface with the stalks, reaching an averaging surface density of 55,000 +/- 10,000 probes/μm^2^ (see Methods, Fig. S1), ∼10-fold increase compared to those frequently used methods^6^ (Fig. 1D).

Following the substrate fabrication step are the standard protocols for sequencing-based spatial transcriptomics, which can be completed in any biological research lab equipped with a cryo- microtome. The standard protocols include frozen tissue micro-sectioning, tissue slice being mounted on the substrate, fixation, permeabilization, and mRNA capturing by the probes. Subsequently, reverse transcription and second-strand synthesis are performed. The second strands are then denatured, washed off, and collected. The resulting cDNAs are utilized for library construction and subjected to paired-end sequencing in conjunction with SBCs. Through the alignment of each cDNA to the SBC map, a high-resolution spatial transcriptomic landscape is constructed.

### Spatial cellular-type architecture of mouse testis

We applied Salus-STS to a mouse testis tissue section and achieved high sensitivity of mRNA detection with a median of 13,834 UMIs and 4,199 genes per 10 × 10 μm² bin at a saturation level of 0.43. The UMI and gene numbers are significantly higher than those typically reported in scRNA-seq analysis^26^ (Fig. 2A). The spatial distribution of Uniform manifold approximation and projection (UMAP) clusters delineates the architecture of seminiferous tubules and the developmental stages of germ cells (Fig. 2B). We integrated the 10 × 10 μm² bin data from Salus-STS with scRNA-seq data^26^. UMAP analysis demonstrates high consistency between our data and the scRNA-Seq data in terms of cell types and their distributions in the UMAP plot (Fig. 2C). Additionally, the cluster distributions recapitulate the process of germ cell development: from spermatocytes to round and elongating spermatids. These results highlight the capability of Salus- STS to characterize the full transcriptomic landscape.

**Figure 2.**
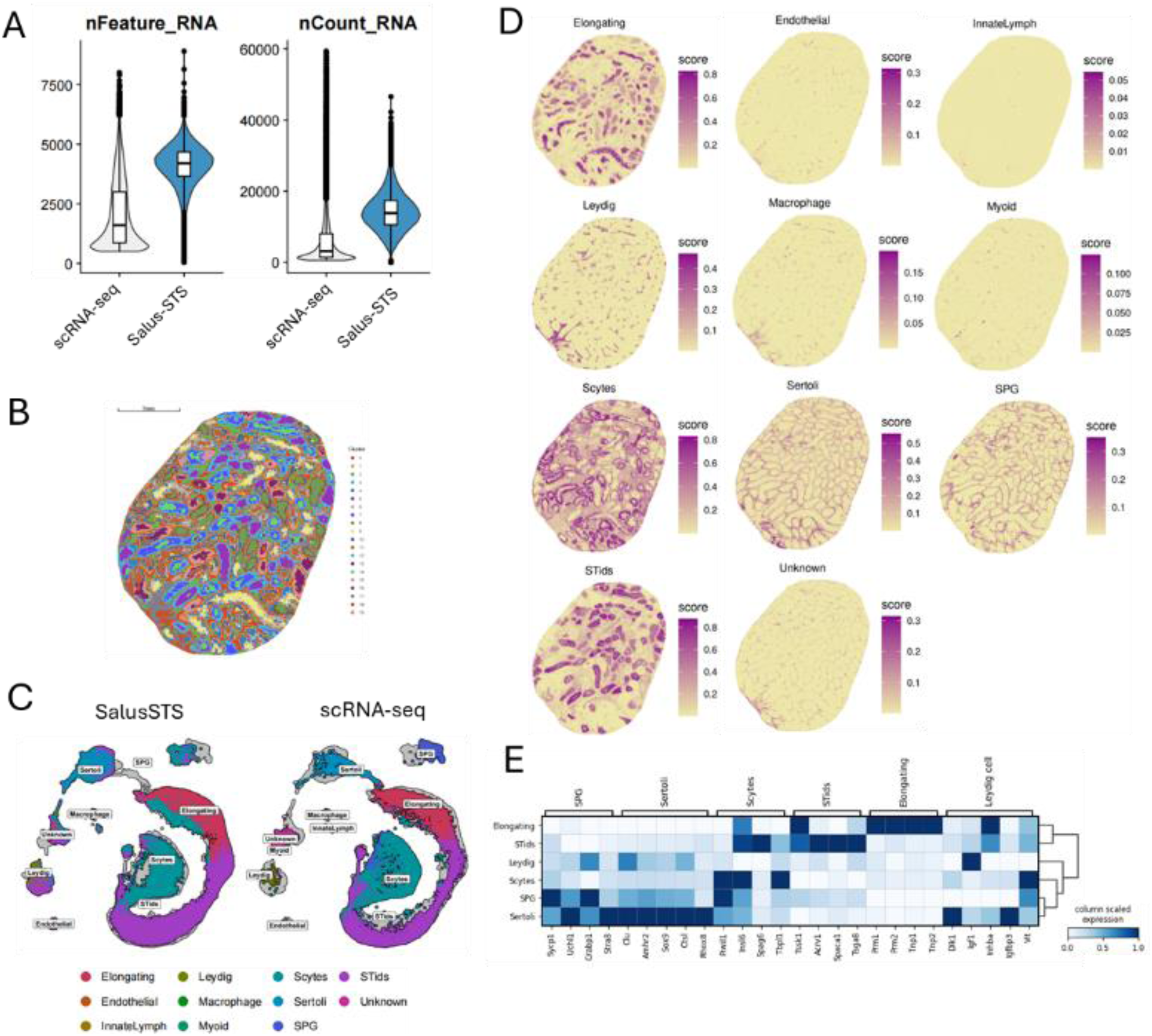
Spatial cellular architecture of mouse testis. (A) Violin plots of number of UMIs and genes detected by Salus-STS (10 × 10 μm^2^ bins) and scRNA-seq (per cell). (B) Spatial mapping of clusters from UMAP analysis. Scale bar: 1mm. (C) UMAP analysis of Salus-STS dataset with scRNA-seq data. SPG: spermatogonia; Scytes: meiotic spermatocytes; Stids: postmeiotic haploid round spermatids; ES: elongating spermatids. (D) Spatial distributions of different cell types annotated by RCTD using 10 ×10 μm^2^ bins. (E) Expression pattern of specific marker genes across major cell types from mouse testis.

To annotate the cell types of the testis tissue, we used RCTD method and single cell data from Gene Expression Omnibus (GEO). Major cell types were properly annotated and mapped (Fig.2D and Fig. S2). Cell annotation was further validated by examining the expression patterns of marker genes across different clusters (Fig. 2E).

Some observations are consistent with the literature and some are new. For example, it has been reported that seminiferous tubules are constructed by tight junctions of Sertoli cells^27^, which are located in the outermost layer of the tubules, consistent with our results. Within these tubules, we observed the different distributions of spermatogonia, spermatocytes, elongating spermatids, and round spermatids in the tubules, which align with our current understanding of mouse testis anatomy^28^.

Additionally, our dataset identifies a broad range of somatic cells, including Leydig cells, endothelial cells, peritubular myoid cells, and macrophages (Fig.2D and Fig. S2). Peritubular myoid cells, which surround the seminiferous tubules, are typically found to be a single layer in rodent testes^29^. Due to their rarity, these cells are often underrepresented in scRNA-seq data^26^. Remarkably, our dataset clearly reveals the structure of peritubular myoid cells, exhibiting a latticework pattern (Fig.2D and Fig. S2). Such a structure has not been reported in the literature using similar methods, demonstrating that the spatial resolution and capture efficiency are crucial in locating rare cells and revealing their spatial organizations.

Interestingly, in another area, we found macrophages and Leydig cells assembled and colocalized (lower left corner of Fig.2D and Fig. 24), together with enriched lymphocytes while sperm cells were absent. This possibly suggests a local inflammation incidence. It has been reported that macrophages and Leydig cells are functionally related and macrophages may produce cytokines that tune the steroidogenesis of the Leydig cell^30,31^. However, how this happens through a variety of signaling pathways is still largely unclear. High resolution, high capture efficiency spatial transcriptomics may suggest clues in solving these problems.

### Analysis of mouse testis architecture based on sub-cellular 2 × 2 μm^2^ bins

Given the large number of genes and UMIs captured, we analyzed the mouse testis dataset at an unprecedented 2 μm resolution. A ∼0.7 × 0.7 mm² area was selected, encompassing approximately seven seminiferous tubules. Using 2 × 2 μm² bins, we identified a median of 215 genes and 333 UMIs (Fig. 3A).

**Figure 3.**
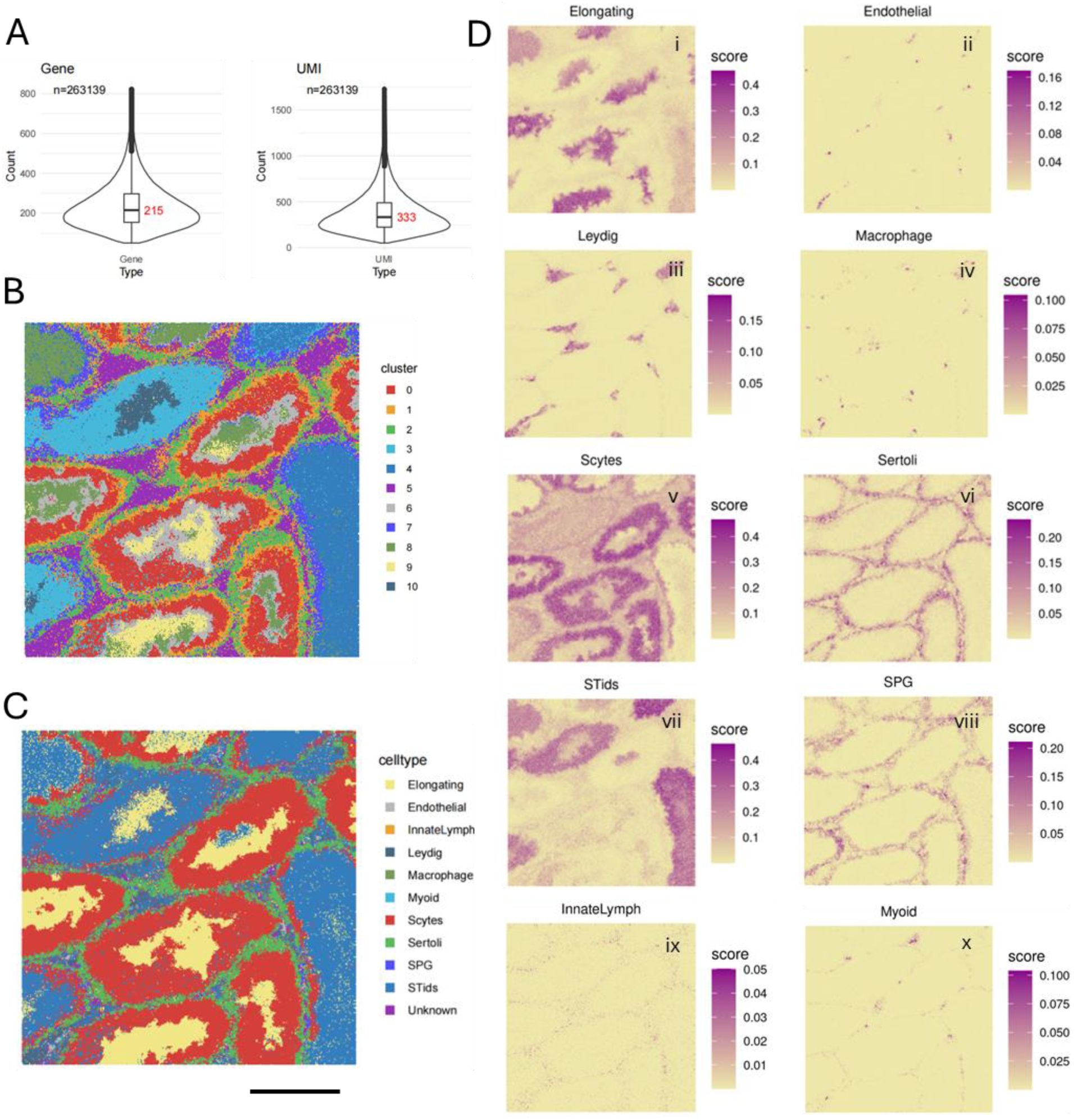
Analysis of mouse testis architecture based on sub-cellular 2 × 2 μm^2^ bins. (A) Violin plots for numbers of UMIs and genes detected per bin. (B) UMAP clustering analysis for all the sub-cellular bins. (C) Spatial distribution of cell types for each bin annotated with RCTD analysis. Scale bar: 200 μm. (D) Spatial distributions of different cell types for each bin.

Unsupervised clustering analysis reveals the seminiferous tubules and layered structures inside the tubules (Fig. 3B), possibly suggesting that these layers are groups of cells belonging to the same types or sub-types. It also indicates that the inter-cell type difference is more significant than the difference between portions of the same cell.

We further annotated each 2 × 2 μm^2^ bin using RCTD method (Fig. 3C). The tissue architecture is very similar to that disclosed by unsupervised clustering. This indicates that both methods are reasonable and valid. The spatial distributions of single cell types revealed more details about the tissue structure (Fig. 3D), allowing us to observe islands of bins with sizes similar to a cell. For example, in Fig. 3C-iv, we see islands of bins which are likely individual macrophages. It has been reported that the transcriptome of Sertoli cells is closely associated with stages of seminiferous epithelial cycles^32,33^. Our spatial architecture demonstrates high resolution Sertoli cell distribution that different Sertoli cells from adjacent seminiferous tubules can be well-separated (Fig. 3C-vi). The high-resolution cell maps undoubtedly will facilitate the study of Sertoli cell-germ cell interactions.

Above analysis based on 2 × 2 μm^2^ bins demonstrates the great spatial resolution Salus-STS has. Note that RCTD, a deconvolution algorithm resolving cell compositions within "spots" containing multiple cells with mixed types, may not be ideal for the analysis with subcellular sized elements. In fact, it is unclear to us so far how to best use the high resolution, high sensitivity data. However, current reasonable results and new structures disclosed suggest that subcellular level spatial transcriptomics is possible. It invites new ideas to dig out rich information buried in these data.

### Spatially resolved whole transcriptome of mouse brain

To illustrate the *in situ* capturing ability of Salus-STS, we analyzed an adult mouse hemibrain tissue section. The spatial heatmaps of UMIs and genes reveal the anatomical structure of the mouse brain section (Fig. 4A). Our dataset achieved a high detection sensitivity for mRNAs, with a median of 2,851 UMIs and 1,340 genes at a saturation of 0.46 for 10 × 10 μm² bins (Fig. 4B), significantly higher than previous reports^13–15^. For examples, Nova-ST reported 349 UMIs and 199 genes for 10 × 10 μm² bins (no saturation reported) for adult mouse brain. For StereoSeq, the original manuscript did not report these values; however, multiple researchers calculated from the published dataset that the UMIs for StereoSeq are between 300∼500 per 10 × 10 μm² bin at a saturation of > 0.9. We confirmed above calculations and concluded that current sequencing-based methods usually detect UMIs in lower hundreds per10 ×10 μm² bin for adult mouse brain, which is significantly lower than that of Salus-STS.

**Figure 4.**
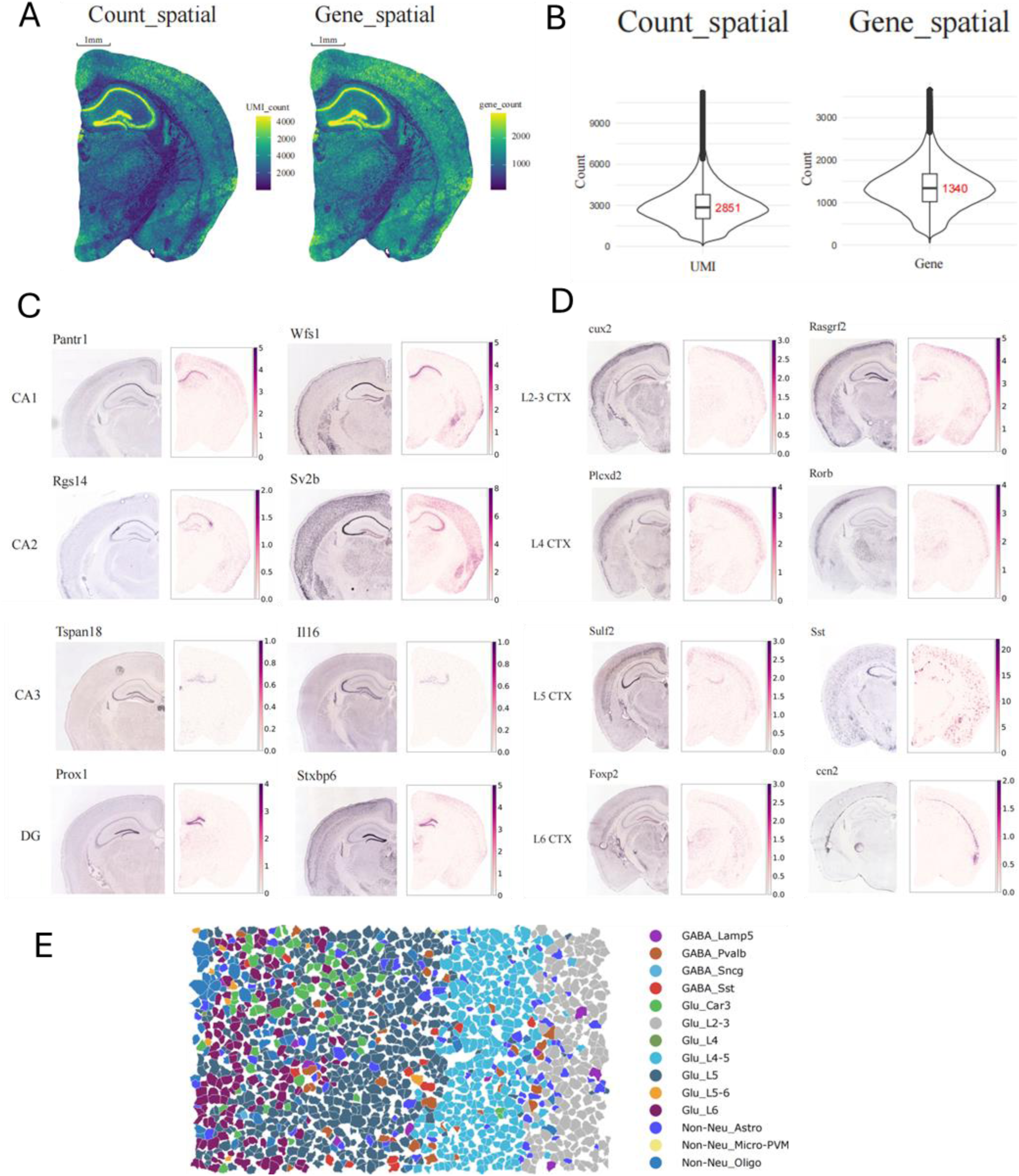
Spatially resolved whole transcriptome of mouse brain. (A) Spatial heatmap of UMIs and genes captured across the mouse brain section. (B) Violin plots of UMIs and genes detected. (C) Spatial distributions of hippocampal markers detected by Salus-STS (right), compared with ISH images (left). (D) Spatial distributions of specific cortical layer markers detected by Salus-STS and compared with ISH images. (E) Spatial distribution of different cell types in the cortical region annotated by RCTD using cellbins.

To showcase the high-definition transcriptome map disclosed by the data, we visualized the spatial distributions of specific marker genes in the hippocampus and cortical areas (Fig. 4C): *Pantr1,* and *Wfs1* for CA1, *Rgs14* and *Sv2b* for CA2, *Tspan18* and *Il16* for CA3, *Prox1* and *Stxbp6* for DG. Additionally, a series of cortical markers exhibit layer-specific distribution patterns (Fig. 4D). Impressively, the distribution of *Ccn2* clearly reveals the structure of L6b sublayer. Overall, we observed a consistent correspondence between the marker distributions and their respective anatomical regions, closely resembling the ISH data from the Allen Brain Atlas^34^. To conclude, using 10 × 10 μm² bins as the elements of analysis and RCTD annotations with a scRNA-seq dataset as the reference^35^, we precisely reconstructed the cellular map of a mouse brain section, identifying 29 subclasses of glutamatergic neurons (Fig. S3), 6 sub-classes of GABAergic neurons (Fig. S4A), and 6 subclasses of non-neuronal cells (Fig. S4B). 41 out of 42 subclasses in the reference dataset were successfully characterized, demonstrating the ability of Salus-STS to identify major and rare cell sub-types.

The mouse cortical area comprises diverse neurons and non-neuronal cells of various types and sizes. To better visualize the cellular landscape, we selected an area from the cortex and performed cell segmentations (i.e., cellbins, see Next section). RCTD annotations successfully mapped 14 distinct cell types, including 7 glutamatergic neuron subtypes (Car3, L2/3, L4, L4/5, L5, L5/6, L6), which shows layer-specific distributions, 4 GABAergic neuron subtypes (Lamp5, Pvalb, Sncg, SST), and 3 non-neuronal types (astrocytes, microglia, and oligodendrocytes) (Fig. 4E). Our dataset clearly reveals the spatial relationships between neurons and non-neuronal cells, which are crucial in cell-cell interactions and communications, such as juxtacrine and paracrine signaling.

### Mouse brain analysis using Salus Cellbins

Fig. 5 shows another example of mouse brains, whose anatomic structure includes the cortex, hippocampus and thalamus, can be accurately identified using unsupervised clustering of 25 × 25 μm^2^ bins (Fig. 5A). In this case, we analyzed the hemibrain using our cell segmentation algorithm (Salus Cellbins), which recognizes cell nuclei based on unspliced mRNA distribution and employs a watershed algorithm to define cell boundaries (Fig. 5B).

**Figure 5.**
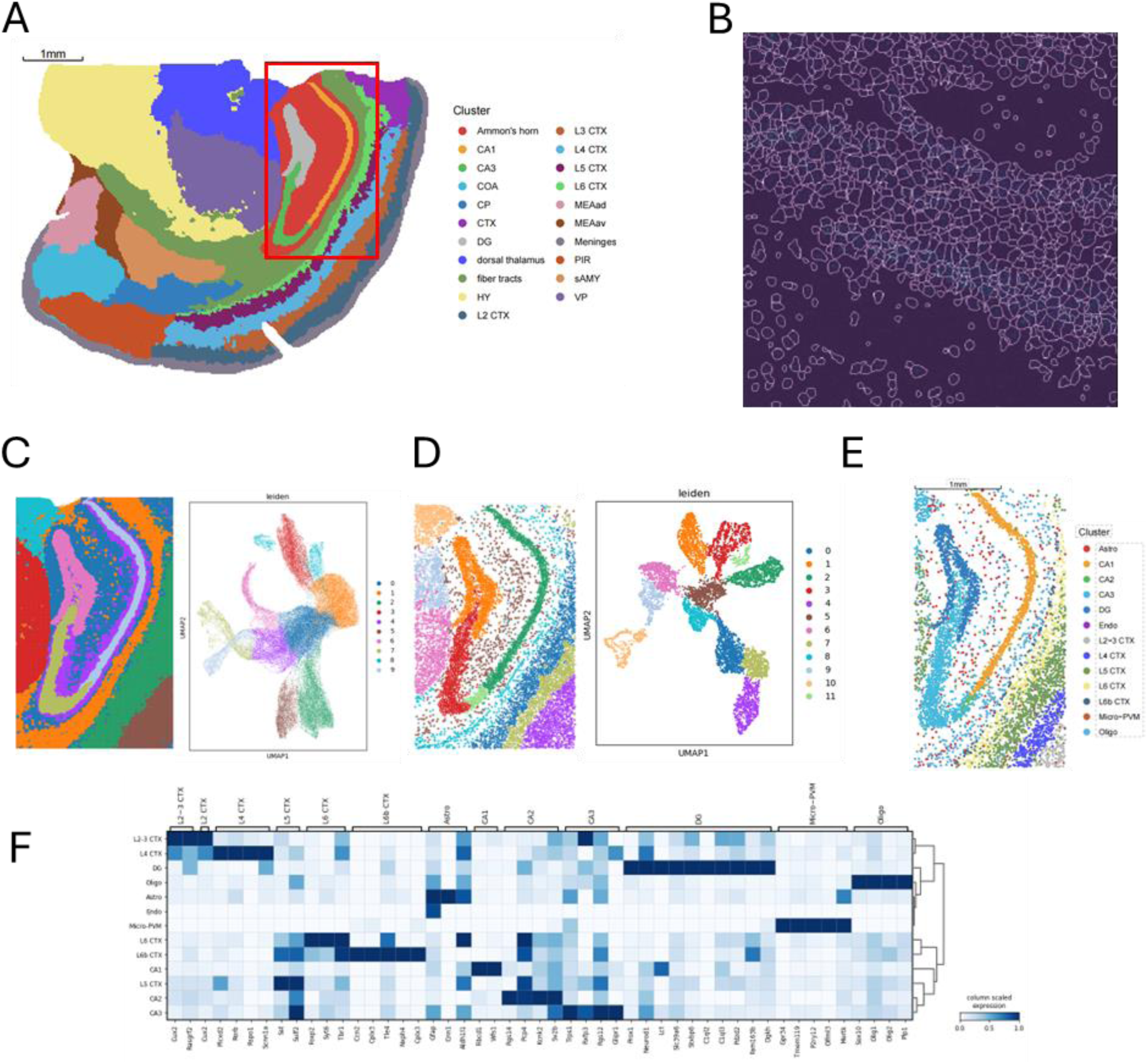
Whole transcriptome of mouse brain analyzed with Salus Cellbins. (A) Unsupervised clustering of the mouse hemibrain section using 25 × 25 μm^2^ bins. Bins are colored by their annotation. CA1: cornu ammonis; COA: cortical amygdalar area; CP: caudoputamen; CTX: cortex; DG: dentate gyrus; HY: hypothalamus; MEAad: medial amygdalra nucleus, anteroventral part; MEAav: medial amygdalar nucleus, anterodorsal part; PIR: pifiform area; sAMY: stiatum-like amygdalar nuclei; VP: ventral posterior complex. (B) Visualization of segmented cell boundaries. (C) UMAP plots of 25 × 25 μm^2^ bins and spatial mapping of the clusters in the red frame from (A). (D) UMAP plots of segmented cells and spatial mapping of the clusters. (E) Segmented cells annotated by RCTD. (F) Expression pattern of specific marker genes across different cell types from hippocampus.

Notably, cell segmentation significantly improved clustering quality in the hippocampus area by clearly distinguishing different regions as compared to the original 25 μm ×25 μm^2^ bin result (Fig. 5C, D). It has been reported CA2 area resembles a terminal portion of CA3 region^36^, thus is often recognized as a part of CA3. However, CA2 exhibits distinct molecular and functional properties, as well as unique connectivity patterns that may be relevant to disease mechanisms^37^. Leveraging cell segmentation and annotation, Salus-STS successfully resolved the hippocampal subdivisions, including all DG, CA1, CA2, and CA3 (Fig. 5D) regions.

Furthermore, annotations of segmented cells by RCTD^38^ revealed a high-definition cellular landscape in the mouse hippocampus, including clearly separated cortical layers (Fig. 5E). The cell annotation was rigorously verified by comparing the expression patterns of known cell markers across different cell types (Fig. 5F).

### Lateral molecular diffusion

The spatial accuracy of mRNA capture is a critical aspect of spatial transcriptomic methods, typically quantified by the lateral molecular diffusion of mRNAs with known expression patterns. To evaluate this, we profiled the spatial transcriptome of mouse olfactory bulb (MOB). Unsupervised clustering with UMAP at a bin size of 10 × 10 μm^2^ was performed. Clusters were annotated based on expression profiles of specific markers, which aligns well with tissue anatomy from the Allen Brain Atlas^34^ (Fig. 6A).

**Figure 6.**
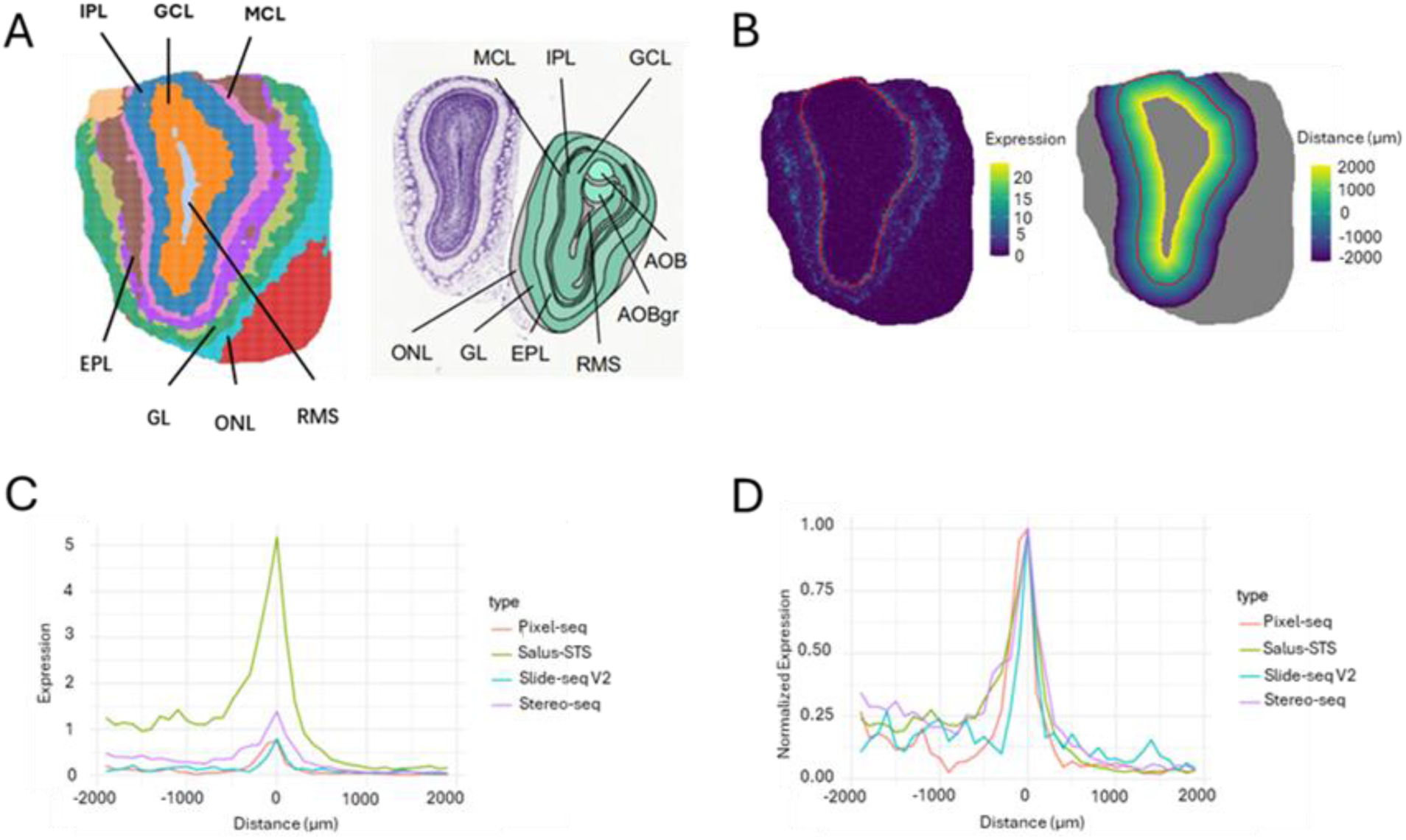
Lateral molecular diffusion analysis. (A) Unsupervised clustering and annotations of mouse olfactory bulb tissue section processed by Salus-STS (left). Refence of mouse olfactory bulb anatomy from Allen brain atlas (right). (B) Selected region for diffusion analysis, with the red line indicating *Slc17ac7* expression peak (left), yellow to purple region representing the selected 4-mm width region, and the color gradient representing the distance from the red line (right). (C) Mean detection of *Slc17ac7* across the selected region. (D) Normalized mean detection of *Slc17ac7* across the selected region.

To evaluate molecular diffusion, we used the spatial distribution pattern of *Slc17ac7* gene, known for its specific expression in Mitral and Tufted cells^39^, as well as in glutamatergic neurons^40^. We focused on its expression in Mitral and Tufted cells by selecting a 4-mm wide region where *Slc17ac7* expression peaked in the middle (Fig. 6B). We then calculated the mean detection level at every 10 μm interval. The resulting mean detection plot demonstrated that Salus-STS offers significantly more sensitive molecular detection compared to the other methods (Fig. 6C). In addition, Salus-STS maintains excellent control over diffusion as compared to the other methods indicated in the normalized detection plot (Fig. 6D).

## DISCUSSION

Spatial transcriptomic methods have emerged as a powerful tool in biomedical research^41–43^. Despite of the technological advancements over the past decade, there still lacks a method that offers simultaneous high resolution, sensitive gene detection, and whole transcriptomic profiling. Some recent sequencing-based methods give a spatial resolution for capture addresses between 0.5 ∼ 2 μm, well below the size of a biological cell. However, due to poor detection efficiency, the data have to be binned to have a size similar to or larger than a cell for further analysis. The advantage of subcellular address resolution cannot be fully exploited. Even after binning, the total number of genes and UMIs that are captured in a voxel equivalent in size to a cell are ∼one order of magnitude smaller than those from single cell RNA sequencing. This introduces inaccuracy and uncertainty in subsequent data analysis.

In this study, we introduce Salus-STS, a sequencing-based method that relies on solid-phase capture arrays. Salus-STS has three key features: (1) high spatial resolution for capture addresses of ∼1 μm; (2) ultra-high probe density of ∼55,000 +/- 10,000 probes/μm^2^, enabling efficient whole transcriptome capture and characterization; (3) flexible tuning of the molecular recognition portions of the capture probes, providing versatility for studying different types of targets.

In studying adult mouse testis and brain, Salus-STS demonstrated the highest sensitivity in the literature to the best of our knowledge. It captured a median of 13,834 UMIs and 4,199 genes per 10 × 10 μm² bin at a saturation of 0.43 for mouse testis, and a median of 2,851 UMIs and 1,340 genes at a saturation of 0.46 for adult mouse brain. These numbers are ∼one order of magnitude higher than literature reports^13–15^, making Salus-STS approaching scRNA-seq in detection sensitivity.

The high detection efficiency allowed us to analyze gene expression profiles for mouse testis with unprecedented 2 × 2 μm^2^ bins, elements much smaller than a biological cell. Reasonable and new results about tissue architecture were obtained, which demonstrates that Salus-STS has great potential pushing sequencing-based spatial transcriptomics toward real sub-cellular resolutions^44^. In summary, we envision Salus-STS as a powerful tool that provides both high spatial resolution and high throughput for advancing research in life sciences and translational medicines.

## Supplementary Information

Four supplementary figures are in the Supplementary Materials.

## Data availability

Testis single cell RNA data at the NCBI under GEO accession number GSE112393. Mouse brain reference single cell data generated by Allen Institute for Brain Science are available at web portal (https://portal.brain-map.org/atlases-and-data/rnaseq). SalusSTS gene expression matrix are available at the Google Drive:

(https://drive.google.com/drive/folders/10krNQShSm7E3bgoM cqxBspR- YSqihr?usp=sharing). OpenST adult mouse hippocampus are available at web portal (https://rajewsky-lab.github.io/openst/examples/getting_started/). Stereo-seq mouse brain at CNGBdb under experiment ID (CNX0422300).

## Acknowledgements

This work is supported by Shenzhen Science and Technology Program “KJZD20230923114220041”.

## Author contribution

GW, EL, LZ, YB, and RL conceived the project. GW, EL, DFu, and RL supervised the whole project and designed the experiments. RL, DFu, YZ and DFeng designed the Salus-STS substrates and cassettes. XZ, YiC, JC, and XL performed the majority of the experiments. ZX and HX designed the bioinformatic pipeline. HL, RG, CL, and ZX analyzed the data. GL participated in designing functional oligos. QC optimized enzyme for the biochemical reactions. WC, YC, LC customized the sequencer for spatial transcriptomics study. RL, ZX, DFu and GW wrote the manuscript. All authors read and approved the manuscript.

## Author information

All authors are employees of Salus BioMed Inc. Ltd. Correspondence and requests for materials should be addressed to.

## METHODS

### Animal Tissue Samples

All relevant procedures involving animal experiments presented in this study are compliant with ethical regulations regarding animal research and were conducted under the approval of the Animal Care and Use committee of the Guangzhou Institutes of Biomedicine and Health, Chinese Academy of Sciences (license number TOPGM-IACUC-2024-0255). Male C57BL/6J mice (4-5 weeks old, Top Biotech, Shenzhen, China) were sacrificed, mouses organs were immediately dissected. After collection, tissues were embedded in optimal cutting temperature compound and snap-frozen on dry ice, stored at -80 ℃ before cryo-sectioning.

### Preparation of Salus-STS Capture Substrates

We synthesized the barcode library (Genescript), and the poly-T probe library. The barcode library carries a 30-nt random nucleotide sequence that serves as a spatial barcode (SBC). The probe library contains a 10-nt random nucleotide sequence as the unique molecular identifier (UMI).

The barcode library was loaded onto the capture substrate that was modified from a DNA sequencing chip (Salus Biomed, Shenzhen, China) according to the Salus Pro^TM^ sequencer manual. Bridge amplification was then carried out in the Salus Pro^TM^ sequencer to generate the DNA clusters. To decipher the SBC of each cluster, the SBC primer was used and a single-ended 30bp sequencing was processed in the Salus Pro^TM^ sequencer. After sequencing, a second bridge amplification was processed to fill the gaps between clusters. An enzymatic cleavage was then performed to remove the reverse strands. To generate the capture probe, the probe library was then hybridized to perform an extension reaction (1 μM probe library in Kapa Hifi HotStart ReadyMix). The probe library was then washed off through denaturation.

### Calculation of probe density on Salus-STS slide

We first fabricated slides modified with poly-T oligos of different density. The slides were then hybridized with cy5-labeled poly-A oligos and imaged to quantify the fluorescent intensity. We plotted the standard curve of fluorescent intensity vs probe density. To characterize the probe density of Salus-STS slide, we hybridized the slide with Cy5-labeled poly-A oligos, the probe density was calculated from the fluorescent intensity using the standard curve.

### Salus-STS library preparation

#### Tissue processing, H&E staining and imaging

Tissues were sectioned at 10-μm thickness in a Leica CM1950 cryostat and mounted on the Salus- STS chip. The chip was immediately baked at 37 ℃ on a slide dryer for 3 min. Then, the chip was submerged in pre-chilled methanol and fixed for 30 min at -20 ℃.

H&E staining was performed by incubation with hematoxylin for 5 min, bluing buffer for 2 min, and eosin for 2 min. Tissue sections were scanned on a Nikon Ni-U microscope under brightfield.

#### Permeabilization, reverse transcription, and second strand synthesis

Tissue sections were permeabilized in 0.1% pepsin in 0.1 M HCl buffer, at 37 ℃ for 3 min, then washed in 0.01X SSC with 5% RNase inhibitor. Reverse transcription was performed by incubation of sections in RT solution (10U/μL Maxima H Minus Reverse transcriptase, 1mM dNTPs, 5% RNase inhibitor in 1X Maxima RT buffer) at 42 ℃ for 3 h, followed by a wash with 0.01X SSC with 5% RNase Inhibitor). The sections were then treated with Exonuclease I digestion (2 U/μL Exo I enzyme in 1X Exo I buffer, 37 ℃, 1 h) and washed with 0.01X SSC. After that, 80 mM KOH was used to digest the tissues by incubation at room temperature for 15 min, then neutralized by buffer EB (10 mM Tris-Cl, pH 8.5).

The second strand synthesis mix (0.5 U/μL Klenow Fragment (3’-->5’ exo-), 0.5 mM dNTPs, 5 μM random primer in 1X NEBuffer 2.0) was prepared and the sections were incubated for 1 h at 37 ℃. Then the sections were washed with buffer EB.

#### Library preparation and sequencing from Salus-STS

To collect the cDNA library, the cDNA was denatured by 80 mM KOH and neutralized with (1M Tris-HCl, pH 7.0). The cDNA library was first amplified with 2.5 μM cDNA primers in Kapa Hifi HotStart ReadyMix). PCR reaction: 95 ℃ 3 min, 15 cycles of (95 ℃ 30 s, 60 ℃ 1 min, 72 ℃1 min), 72 ℃ 2 min and 4 ℃ infinite.

50 ng cDNA product was then fragmented with in-house Tn5 transposase at 55 ℃ for 10 min, then quenched by 0.02% SDS for 5 min at room temperature. The product was further amplified with 2 μM primers in Kapa Hifi HotStart ReadyMix. PCR reaction: 95 ℃ 5 min, 15 cycles of (95 ℃ 20 s, 60 ℃ 20 s, 72 ℃ 30 s), 72 ℃ 5 min and 4 ℃ infinite.

DNA product was sequenced in the Salus Pro sequencer according to the manufacturer’s instructions, in a pair-ended (100-150bp) mode.

### Data analysis

#### Salus-STS raw data processing

Fastq files were generated from Salus Pro sequencer. SBCs were acquired from 1-30 bp of read 1, while UMIs were acquired at 88-97 bp of read 1. SBCs were first mapped to sequencing results generated while Salus-STS chip preparation, which contains both sequences and coordinates of each SBC. The mapping allows 1-bp mismatch to correct sequencing and PCR errors. Read 2 sequences, which carried the cDNA information, were aligned to the mouse reference genome (mm10) using STAR^45^ and annotated to their corresponding genes. UMIs was used to distinguish distinct mRNA captures with the same SBC and to remove PCR duplicates. An expression profile was generated with coordinates of each transcript.

#### Tissue Boundary Recognition

Tissue boundary recognition was processed by first segment tissue area from H&E images, using Segment Anything Model (SAM, https://segment-anything.com/), generating tissue boundary masks. Then, the mask was applied to spatial transcriptomic UMI heatmap, by custom script, and manual alignment when needed.

#### Unsupervised Clustering

Tissue area was divided into non-overlapping square bins with side length of 10 μm, 25 μm, and 100 μm, then the expression profile matrices were generated based on different bin size. Unsupervised clustering of bins was processed using STAGATE^46^.

#### Cell Segmentation

We employed the watershed algorithm to identify cells from the RNA expression matrix. Specifically, we aimed to delineate concentrations of unspliced RNA as cell cores, given that unspliced RNA fragments tend to be located within the cell nucleus. This property is particularly useful for recognizing cell boundaries and dividing the chip into cell-specific regions. To achieve this, we first applied a Gaussian function to blur the RNA count distribution, as described by the RNA expression matrix, in order to suppress random noise. We then used the watershed algorithm to detect signal concentrations, which were treated as cell core regions. Following this, the region boundaries were extended to capture dissociative signals between cores based on sample conditions and project requirements, taking into account both the size limitation of regions and the total RNA count available for each cell. In some extreme cases where no clear RNA concentrations were observed across the chip, we divided the chip into a series of random regions, each approximately the size of a cell. Before cell segmentation, we used the Segment Anything Model to delineate the tissue-covered area and subsequently identified cells within this region.

#### Cell annotation

Cell annotation was processed by integration of scRNA-seq data with spatial transcriptomic data generated by Salus-STS. Cell2location^47^ or RCTD was used to calculate the relative and absolute abundance of each cell type at each location.

Singe cell data used for testis annotation were accessible through https://www.ncbi.nlm.nih.gov/geo/query/acc.cgi?acc=GSE112393

---

**Fig. S1.**
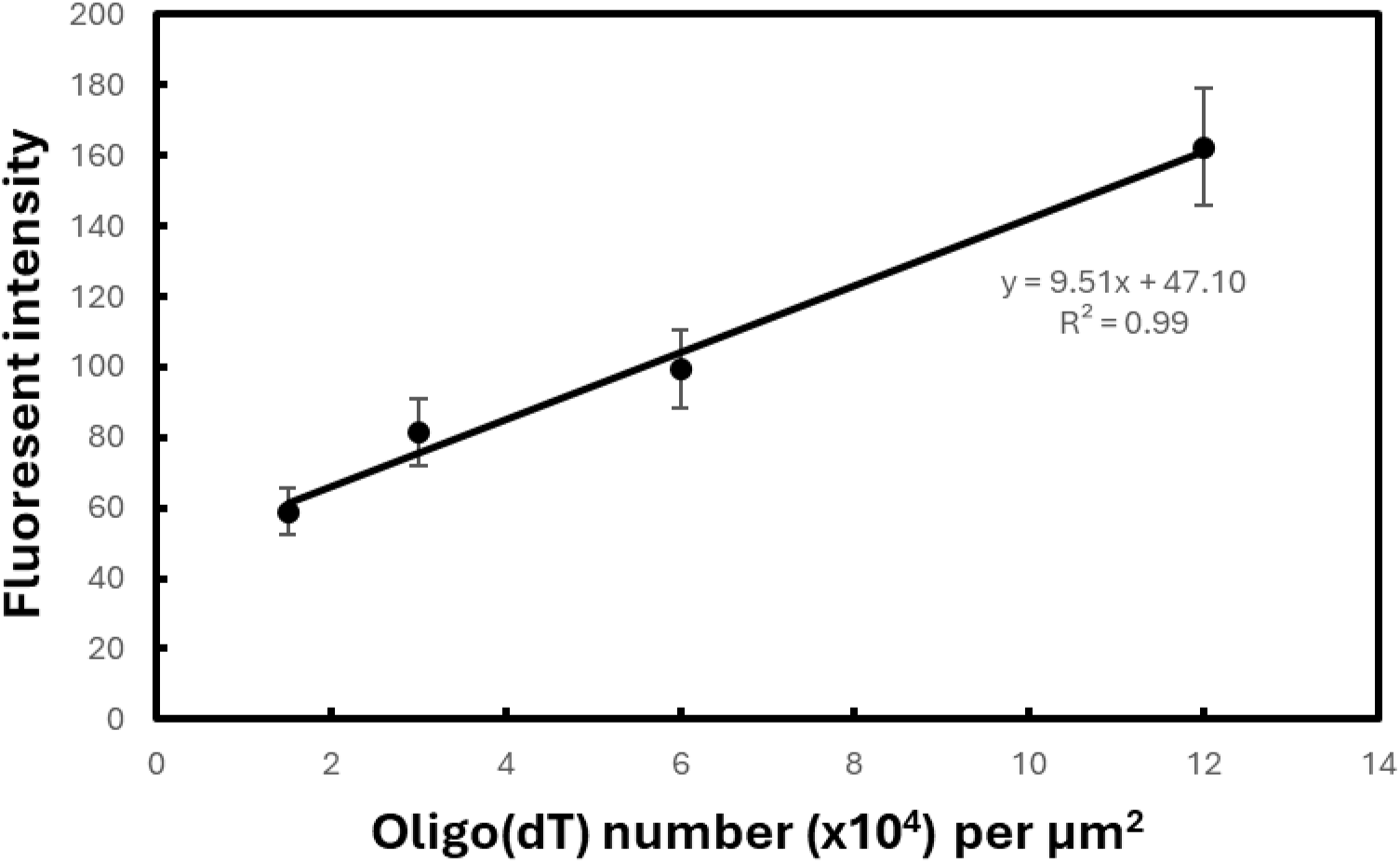
Standard curve of fluorescent intensity of cy5 vs Oligo(dT) number of chip surface area.

**Fig. S2.**
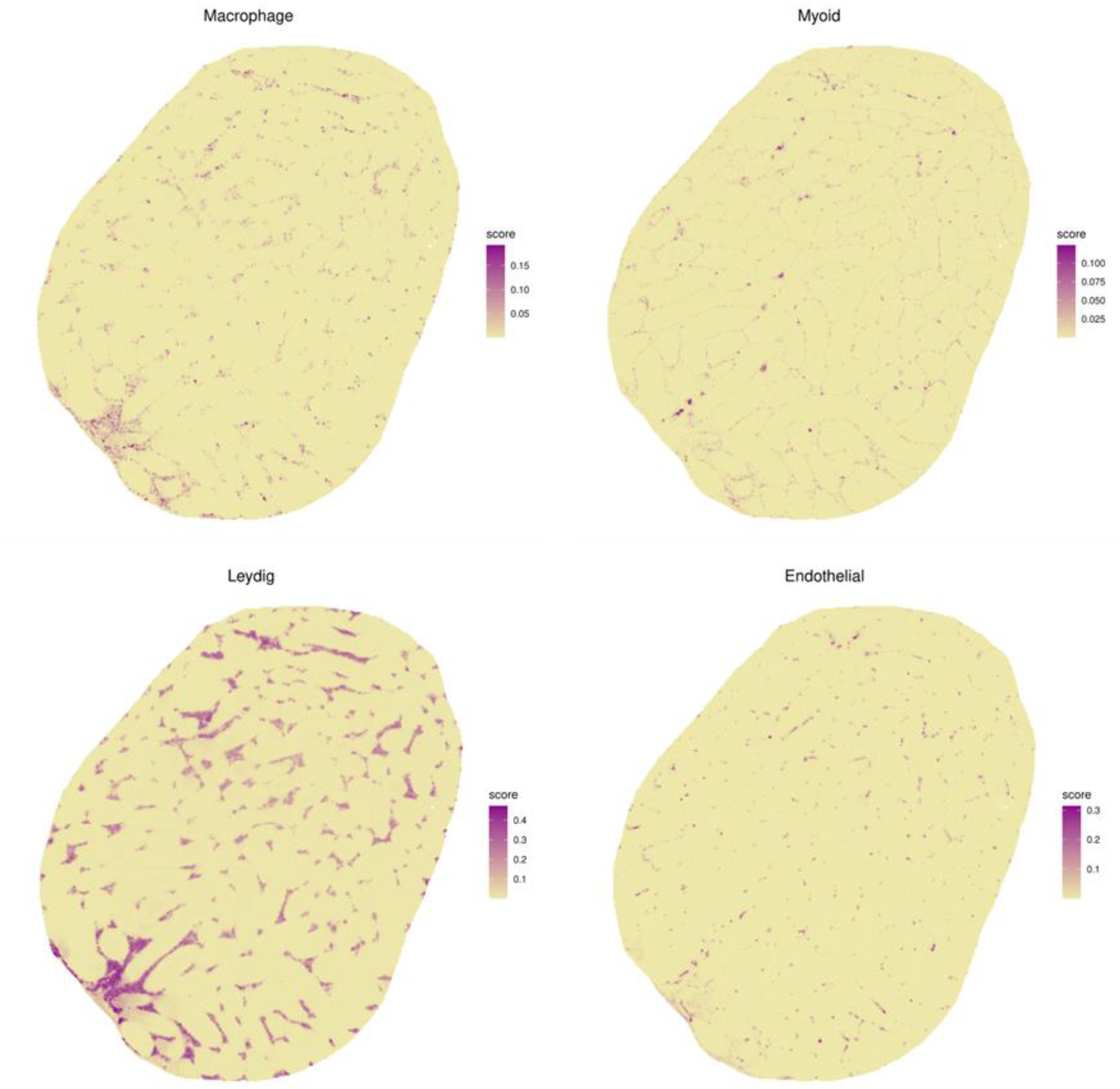
Spatial distributions of different cell types resolved by RCTD.

**Fig. S3.**
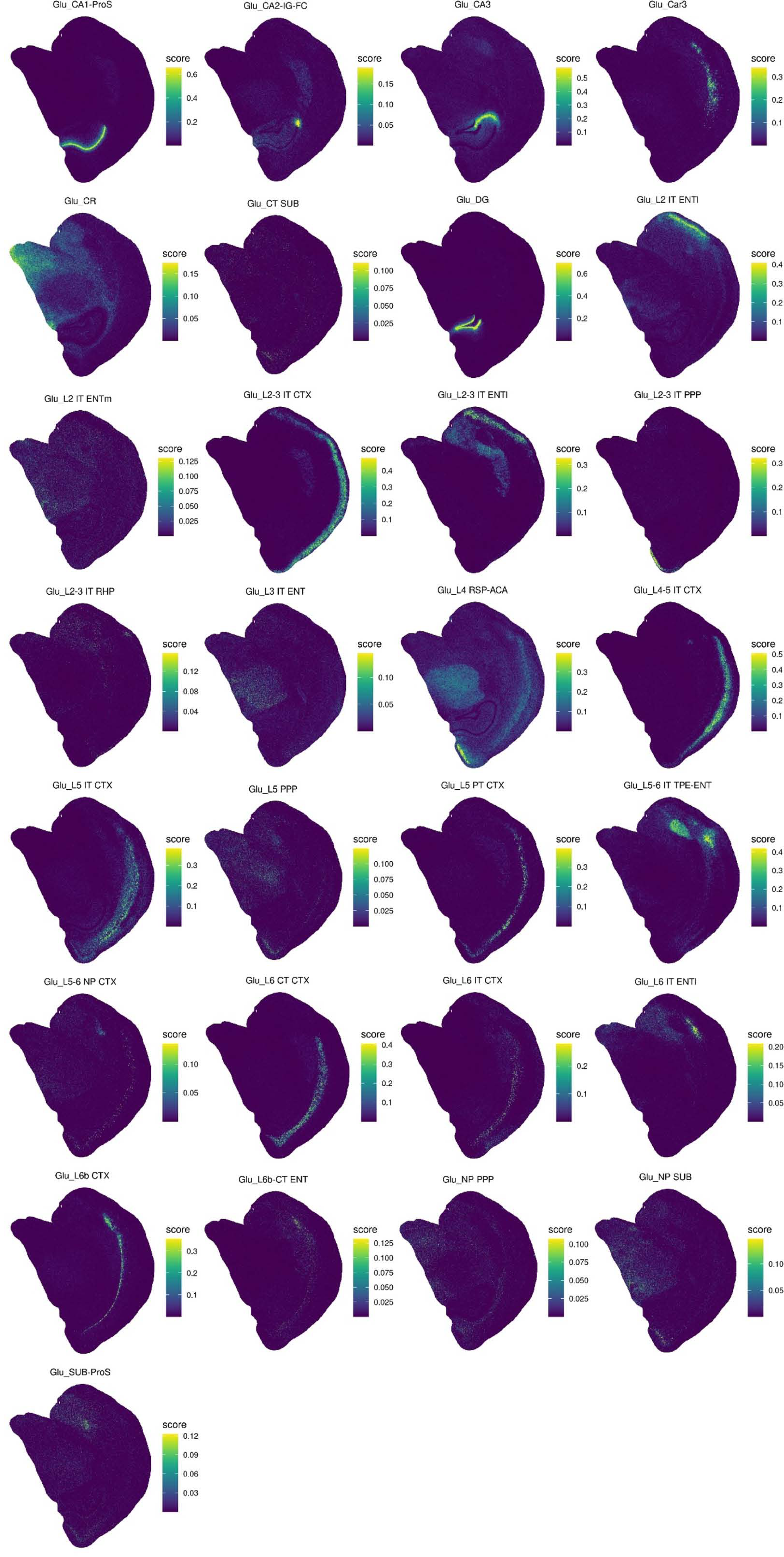
Spatial distributions of RCTD annotated Glutamatergic neurons.

**Fig. S4.**
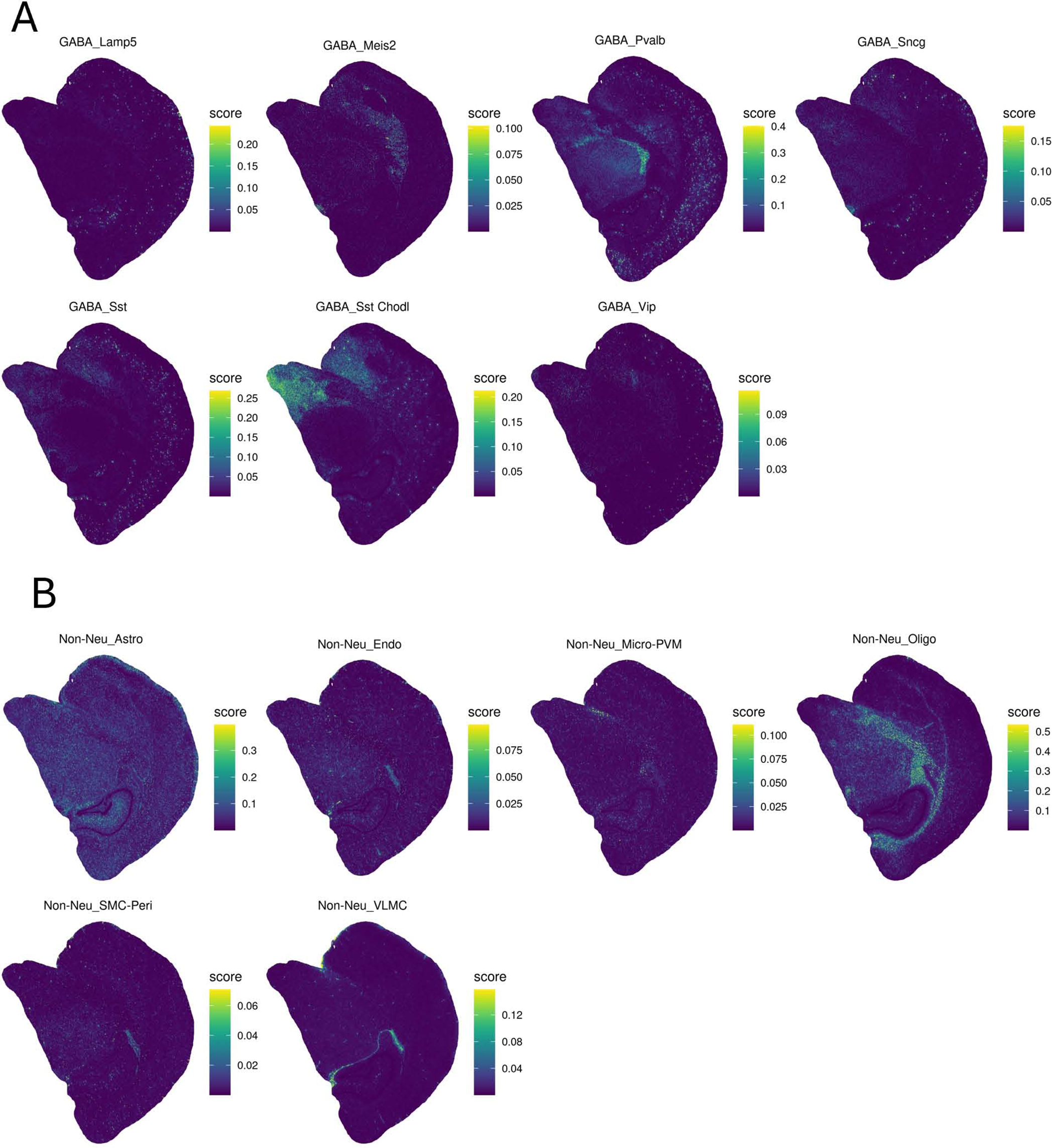
Spatial distributions of RCTD annotated (A) GABAergic neurons and (B) non-neuron cells.

## Notes

### Competing Interest Statement

A patent related to this work has been filed by Salus Biomed.

